# Sharing Data from the Human Tumor Atlas Network through Standards, Infrastructure, and Community Engagement

**DOI:** 10.1101/2024.06.25.598921

**Authors:** Ino de Bruijn, Milen Nikolov, Clarisse Lau, Ashley Clayton, David L Gibbs, Elvira Mitraka, Dar’ya Pozhidayeva, Alex Lash, Selcuk Onur Sumer, Jennifer Altreuter, Kristen Anton, Mialy DeFelice, Xiang Li, Aaron Lisman, William J R Longabaugh, Jeremy Muhlich, Sandro Santagata, Subhiksha Nandakumar, Peter K Sorger, Christine Suver, Nikolaus Schultz, Adam J Taylor, Vésteinn Thorsson, Ethan Cerami, James A Eddy

## Abstract

The Data Coordinating Center (DCC) of the Human Tumor Atlas Network (HTAN) has played a crucial role in enabling the broad sharing and effective utilization of HTAN data within the scientific community. Data from the first phase of HTAN are now available publicly. We describe the diverse datasets and modalities shared, multiple access routes to HTAN assay data and metadata, data standards, technical infrastructure and governance approaches, as well as our approach to sustained community engagement. HTAN data can be accessed via the HTAN Portal, explored in visualization tools—including CellxGene, Minerva, and cBioPortal—and analyzed in the cloud through the NCI Cancer Research Data Commons nodes. We have developed a streamlined infrastructure to ingest and disseminate data by leveraging the Synapse platform. Taken together, the HTAN DCC’s approach demonstrates a successful model for coordinating, standardizing, and disseminating complex cancer research data via multiple resources in the cancer data ecosystem, offering valuable insights for similar consortia, and researchers looking to leverage HTAN data.

## Introduction

The Human Tumor Atlas Network (HTAN) was launched by the National Cancer Institute (NCI) in September 2018, under the umbrella of the U.S. Cancer Moonshot^SM^ program.

The Cancer Moonshot aims to accelerate cancer research and treatment, and has a specific focus on enabling scientific discovery, fostering greater collaboration, and improving the sharing of cancer data^1^. HTAN is a step towards realizing these goals, with a mission to construct three-dimensional atlases of the dynamic cellular, morphological, and molecular features of human cancers as they evolve from precancerous lesions to advanced diseases. As a consortium, HTAN seeks to define critical processes and events throughout the life cycle of human cancers, including the transition of pre-malignant lesions to malignant tumors, the progression of malignant tumors to metastatic cancer, tumor response to therapeutics, and the development of therapeutic resistance. In line with the broader goals of the Cancer Moonshot, HTAN is also committed to rapid and broad sharing of all data with the wider scientific community.

In the broader context of cancer research, HTAN draws upon and extends The Cancer Genome Atlas (TCGA)^2^, a landmark cancer genomics program that molecularly characterized over 11,000 primary tumors and matched normal samples spanning 33 cancer types. TCGA generated comprehensive, multi-dimensional maps of the key genomic changes in major types and subtypes of cancer, providing an invaluable resource for the cancer research community. HTAN is also part of a larger global effort to understand the human body at an unprecedented level of detail. Other initiatives, such as the Human Cell Atlas (HCA)^3^ and the Human BioMolecular Atlas Program (HuBMAP) consortium^4^, are working to create comprehensive, high-resolution maps of all human cell types—healthy and diseased—as a basis for both understanding fundamental human biological processes and diagnosing, monitoring, and treating disease.

While previous large cancer data sharing efforts, such as TCGA, had their own complexities, HTAN presents a new set of challenges. First, each HTAN Atlas is unique and focused on answering different hypotheses regarding cancer progression. As such, HTAN Centers (i.e., U2C awardees responsible for collecting and sharing data related to a particular tumor atlas research program) are free to use whatever experimental assay supports their study. They currently generate a highly diverse set of data types, including bulk sequencing, single-cell sequencing, multiplex imaging, and spatial transcriptomics (**Fig. 1A**). Second, many of the experimental assays used within HTAN—particularly spatial profiling assays—are cutting-edge, and centers are responsible for creating their own bioinformatics pipelines to perform analyses. Third, HTAN is focused on understanding temporal changes in cancer, and the HTAN data model must therefore be capable of capturing longitudinal clinical/phenotype and profiling data. Fourth, the multi-modal nature of HTAN data requires multiple visualization and data access resources, each of which must be tailored to individual data types or end-users.

**Figure 1.**
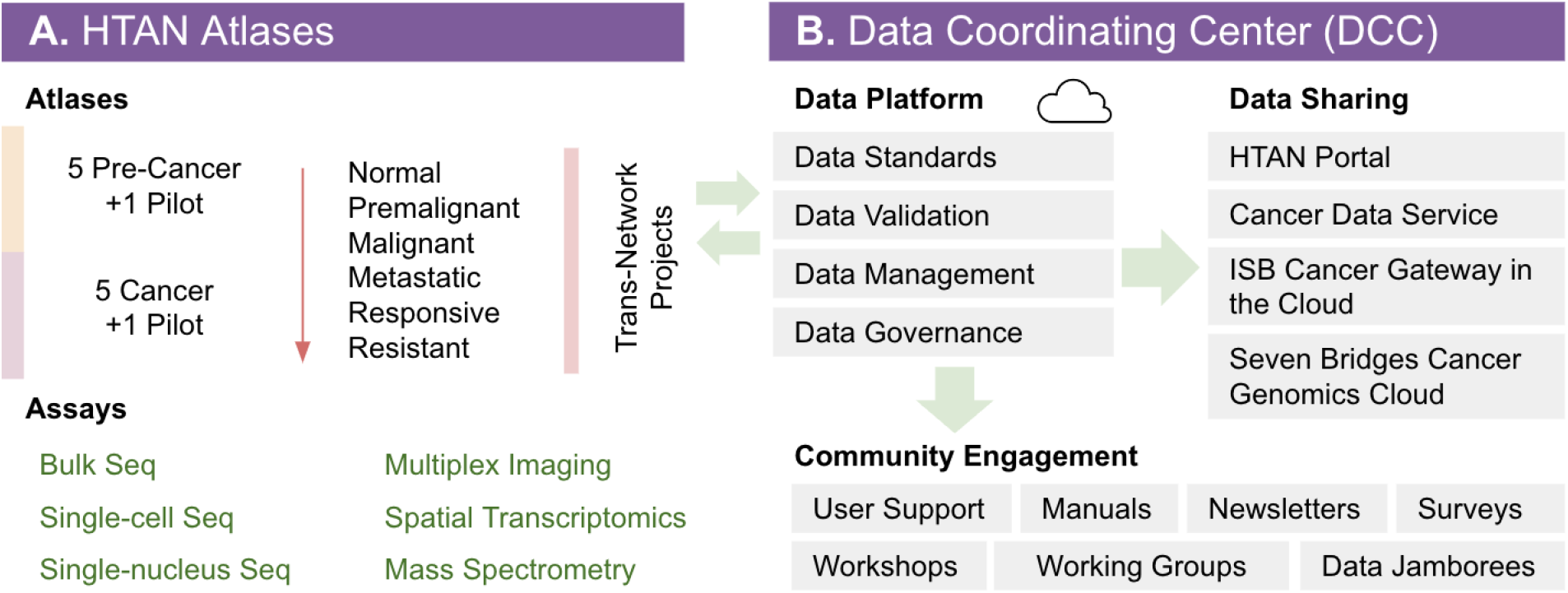
Overview of the HTAN Network and the HTAN Data Coordinating Center (DCC). A) HTAN Atlases focus on specific transitions in cancer and generate a highly diverse set of data types. B) The HTAN DCC is responsible for developing data standards, managing data, and sharing data with the scientific community.

To address the unique challenges of HTAN data, the network includes a dedicated Data Coordinating Center (DCC). The DCC is currently managed by personnel from four institutions: Dana-Farber Cancer Institute, Sage Bionetworks, Memorial Sloan Kettering Cancer Center, and the Institute for Systems Biology. The DCC has overall responsibility for developing HTAN data standards, managing HTAN data within a common cloud infrastructure, and sharing HTAN data with the scientific community (**Fig. 1B**). The DCC infrastructure includes centralized data ingestion, distributed data dissemination, user-friendly portals, and visualization tools. These activities are critical to ensuring that the wealth of data generated by HTAN is available for use by the broader scientific community.

## Results

Launched in September 2018, the first phase of HTAN will be completed in 2024. Here we describe the diverse datasets generated and shared in this phase, the multiple ways users can access HTAN data and metadata, the associated data standards, the enabling technical infrastructure and governance approaches underlying the DCC, and how community engagement is maintained throughout.

### Available Data and Data Levels

HTAN data are now available for two Pilot Projects, ten Atlases, and three Trans-Network Projects (TNPs) (**Table 1**). As of May 2024, this includes 1,945 research participants, 7,833 biospecimens, and profiling data from >20 different molecular assay types. Clinical and biospecimen data are collected and made available in tabular form. Assay data is organized into levels (**Table 2**) similar to prior efforts by the TCGA, with lower levels indicating more raw data and higher levels corresponding to data processing by one or more bioinformatics pipelines; each level for a particular data type adheres to a distinct, standard schema for file formats, metadata fields and values, as well as any additional data validation logic

**Table 1:** HTAN Atlases, organized by Atlas Type and Area of Focus. TNP = Trans-Network Project. More details can be found on the HTAN Portal.

| ID | Lead Institution or Atlas Name | Atlas Type | Area of Focus (Grant/Contract) |
| --- | --- | --- | --- |
| HTA1 | Human Tumor Atlas Pilot Project (HTAPP) | Tumor Atlas | Pilot Project (HHSN261201500003I) |
| HTA2 | Pre-Cancer Atlas Pilot Project (PCAPP) | Pre-Cancer Atlas | Pilot Project (U01CA196[383,386,387,390,403,405,406,408]) |
| HTA3 | Boston University | Pre-Cancer Atlas | Lung (U2CCA233238) |
| HTA4 | Children's Hospital of Philadelphia | Tumor Atlas | Pediatric (U2CCA233285) |
| HTA5 | Dana-Farber Cancer Institute | Tumor Atlas | Multiple Cancer Types (U2CCA233195) |
| HTA6 | Duke University | Pre-Cancer Atlas | Breast (U2CCA233254) |
| HTA7 | Harvard Medical School | Pre-Cancer Atlas | Melanoma, Colorectal Cancer, and Clonal Hematopoiesis (U2CCA233262) |
| HTA8 | Memorial Sloan Kettering Cancer Center | Tumor Atlas | Multiple Cancer Types (U2CCA233284) |
| HTA9 | Oregon Health Science University | Tumor Atlas | Breast (U2CCA233280) |
| HTA10 | Stanford University | Pre-Cancer Atlas | Familial Adenomatous Polyposis (U2CCA233311) |
| HTA11 | Vanderbilt University | Pre-Cancer Atlas | Colorectal (U2CCA233291) |
| HTA12 | Washington University in St. Louis | Tumor Atlas | Multiple Cancer Types (U2CCA233303) |
| HTA13 | Shared Repositories, Data, Analysis and Access (SARDANA) | TNP Atlas | Technology Comparison |
| HTA14 | Tissue MicroArray (TMA) | TNP Atlas | Technology Comparison |
| HTA15 | Standardized Repository of Reference Specimens (SRRS) | TNP Atlas | Technology Comparison |

**Table 2:** Levels of HTAN Data. Lower levels indicate raw data, and higher levels indicate data analyzed by one or more bioinformatics/image processing pipelines. Three primary categories of data are highlighted.

| Level | Single Cell RNA-Seq | Multiplex Imaging | Spatial Transcriptomics |
| --- | --- | --- | --- |
| 1 | Unaligned sequencing reads, usually in the FASTQ file format. | Raw imaging tiles that require preprocessing such as stitching, registration or background subtraction. Typically TIFF or proprietary format | Unaligned sequencing reads, usually in the FASTQ file format. |
| 2 | Aligned sequencing reads, usually in the BAM file format. | Multichannel image. Usually in the OME-TIFF file format, accompanied by a CSV file containing channel metadata. | Aligned sequencing reads, usually in the BAM file format. |
| 3 | Gene expression matrix. For example, a matrix of all cells by all genes, with expression count.<br>Multiple file formats are supported, including CSV, MTX and h5ad. | Segmentation masks denoting nuclei, cytoplasm, whole cells or regions of interest.<br>Multiple file formats are supported although TIFF and OME-TIFF are recommended | Gene expression matrix. For example, a matrix of all cells by all genes, with expression counts.<br>Multiple file formats are supported, including CSV, MTX and h5ad. |
| 4 | Feature matrix. For example, a matrix of cluster assignments or imputed cell types across all sequenced cells.<br>Multiple file formats are supported, including CSV and h5ad. | Feature matrix. For example, a matrix of mean intensity values per cell and channel<br>Multiple file formats are supported, including CSV and h5ad. | Feature matrix. For example, a matrix of cluster assignments or imputed cell types across all sequenced cells.<br>Multiple file formats are supported, including CSV and h5ad. |

### Accessing Data

HTAN data can be accessed via the HTAN Portal as well as several services within the NCI Cancer Research Data Commons (CRDC)^5,6^, such as the Institute for Systems Biology Cancer Gateway in the Cloud (ISB-CGC)^7^, the Cancer Data Service (CDS), and the Seven Bridges Cancer Genomics Cloud (SB-CGC).

#### HTAN Portal

The primary mode of access is the dedicated HTAN Portal available at: https://humantumoratlas.org/ (**Fig. 2A**). The portal enables researchers to explore, access, and download HTAN data via an intuitive user interface. Users can specifically filter HTAN data via a number of criteria, including HTAN Atlas, disease type, assay type, or data level. User-friendly tools for advanced query and visualization of data are also provided. Via the portal, researchers are directed to relevant routes of data access. For open access Level 3 and 4 data, users can directly download data from the Synapse data management platform (RRID:SCR_006307) following easy and free user registration. For controlled-access Level 1 and 2 genomic/transcriptomic data, as well as for Level 2 imaging data, users are directed to data locations with the CDS. The portal also links out to the HTAN Manual for more detailed information regarding the data model, tools, and data repositories.

**Figure 2.**
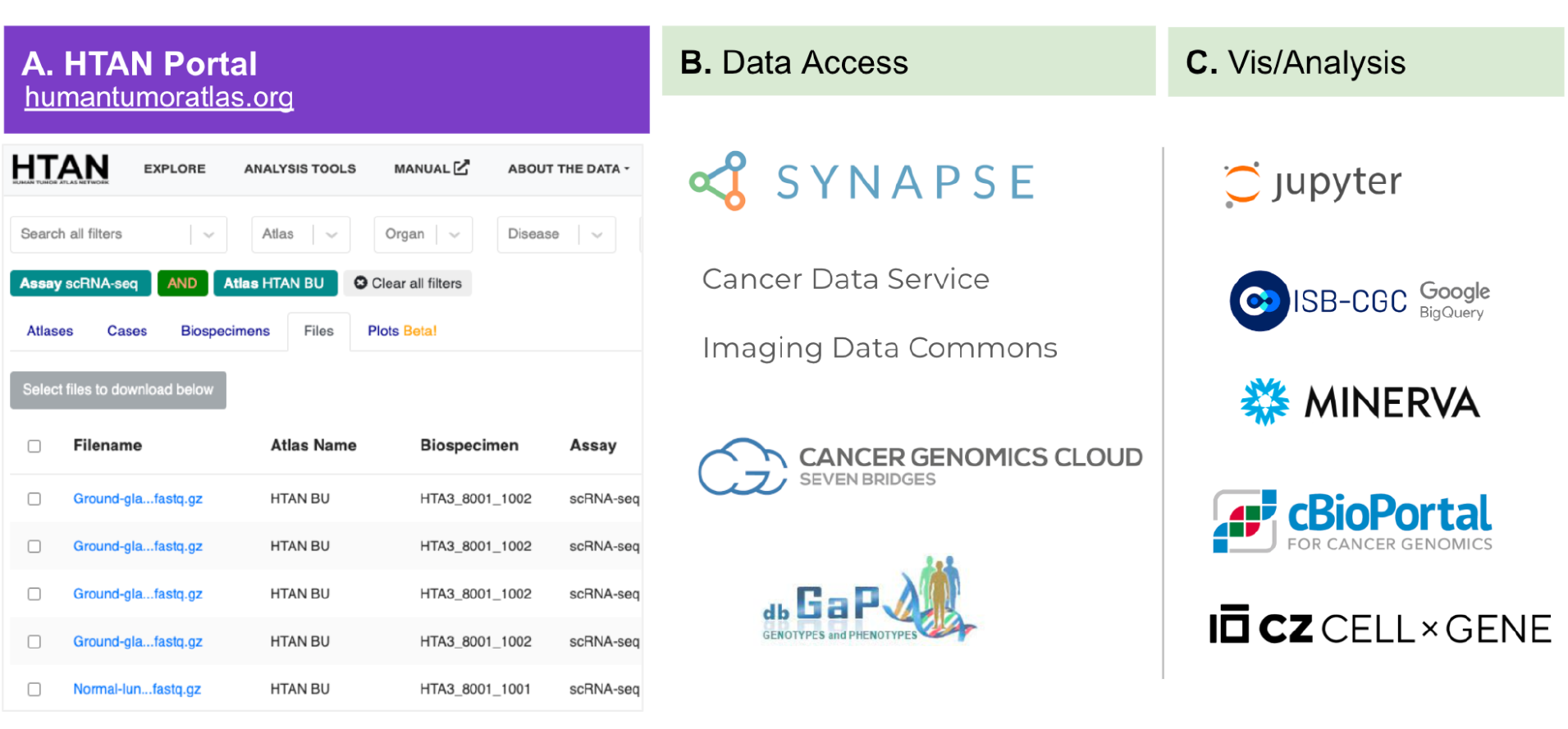
The HTAN Portal provides A) a query interface for finding data and tools, B) data access recipes for lower level 1-2 and higher level 3-4 data, and C) visualization and analysis tools for exploring HTAN data.

#### Visualizing and Analyzing HTAN Data

To enable seamless exploration of HTAN data, the HTAN Portal currently integrates multiple open source visualization and analysis tools (**Fig. 2C**). First, the portal integrates with Minerva, an open source tool developed by Harvard Medical School for visualizing and exploring multiplex imaging data^8^. Two flavors of Minerva are currently supported: (1) Minerva Story, where individual centers expertly annotate and describe specific data sets and delineate specific regions of interest; (2) Auto-Minerva, which auto-generates Minerva images for all multiplex images and assigns reasonable channel defaults for viewing. Second, the portal integrates with cBioPortal for Cancer Genomics, an open source tool for visualizing and analyzing cancer genomics data^9–11^. HTAN datasets with bulk sequencing and other additional methods, including imaging or single-cell sequencing, are deposited into cBioPortal (https://cbioportal.org). Third, the portal integrates with CellxGene, an open source tool developed by the Chan Zuckerberg Initiative (CZI) for visualizing and analyzing single-cell data sets^12,13^. HTAN single-cell data is harmonized for deposition into CellxGene Discover (https://cellxgene.cziscience.com/), enabling exploration of HTAN data with non-HTAN data also in CellxGene Discover.

Finally, HTAN data and metadata are made available in ISB-CGC Google BigQuery. There are numerous BigQuery tables, including metadata tables, single-cell gene expression matrices, and imaging channel data. We also provide numerous example notebooks to illustrate querying and analysis options for HTAN data in ISB-CGC.

#### Controlled-Access Data

For controlled-access Levels 1 or 2 data, users must request access via the NIH database of Genotypes and Phenotypes (dbGaP, Study Accession phs002371). Once approved, users can access HTAN data in the cloud via SB-CGC. The HTAN Portal, ISB-CGC’s Google BigQuery interface, and CDS all provide the functionality to generate Data Repository Service (DRS)^14^ manifest files for seamless access and analysis of HTAN data in SB-CGC. As of May 2024, there are 101 dbGaP-approved data use plans that leverage HTAN data for various innovative applications. For instance, teams integrated HTAN datasets with other genomic datasets to improve the detection of somatic and transcriptional alterations in cancers and aim to identify novel biomarkers for early cancer diagnosis. Similarly, spatial transcriptomics and single-cell RNA sequencing data are being utilized to pinpoint cellular compositions and interactions within tumors, which may reveal new therapeutic targets and strategies. These data reuse projects support the development of predictive models for disease progression and treatment response, ultimately contributing to personalized medicine and improved patient outcomes.

### Data Standards

HTAN has developed a common data model that supports management, standardization, and exploration of clinical, biospecimen, molecular, and imaging data across HTAN Atlases. Clinical data covers demographics, diagnosis, treatment, family history, environmental exposure, and molecular tests. Biospecimen data captures information on storage conditions and provides end-to-end provenance from biopsy to acquired data. Assay metadata (i.e., capturing experimental protocol and instrument context) includes support for bulk and single-cell sequencing, multiplex imaging, and spatial transcriptomics. Complete details are available online at: https://humantumoratlas.org/standards.

The HTAN data model has been generated and is maintained via a community-driven, peer-reviewed process, where members of a working group first assess already established data standards and create a written Request for Comment (RFC) document soliciting community feedback. The RFC documents cover the data itself, and all required and optional metadata elements, and usually undergo several rounds of revision before formal sign-off by all editors. Via this process, the HTAN community has developed a consensus-driven data model that leverages multiple existing data standards and addresses community-driven use cases for data sharing and reuse. The HTAN data model specifically extends the clinical data model developed by the Genomic Data Commons (GDC)^15^, the single cell data model developed by the Human Cell Atlas^3^, and the multiplex imaging model developed by the Minimum Information about Highly Multiplexed Tissue Imaging (MITI) consortium^16^. The data model is continuously evolving and refined based on feedback from the reuse of HTAN data as well as the introduction of novel assays by data submitters.

The HTAN data model is formally represented as an open access and extensible JSON-LD schema document (https://json-ld.org), enabling version control, individual data element links to existing NCI data standards, and the creation of automated validation tools. The JSON-LD schema utilizes the Schema. org specification. In the case of HTAN, this allowed building a data model reusing existing biomedical ontologies when feasible, while adding new HTAN-specific extensions as needed. This promotes interoperability by reusing data elements for experimental variables shared across consortia. It also enhances downstream data discovery via services like Google Datasets Search^17^.

The model comprises 1000+ attributes across 30+ modalities, analysis, and data processing types. A set of 113 HTAN common data elements have been committed to the NCI Cancer Data Standards Registry and Repository (caDSR)^18^, ensuring that these data elements are available to the scientific community through the caDSR portal, API, and tools. These data elements may be collectively browsed and retrieved under the HTAN classification.

### Infrastructure

A broad range of tools, data standards, and platforms were leveraged, enhanced, or developed to support the overall HTAN DCC data infrastructure. This includes tooling to support data and metadata ingestion, data storage, access controls, quality assurance, data sharing, image processing, visualization and analysis (**Table 3**). All data standards and most tools are available via GitHub (https://github.com/ncihtan), and are freely available to other consortia that wish to build upon the work of HTAN.

**Table 3:**
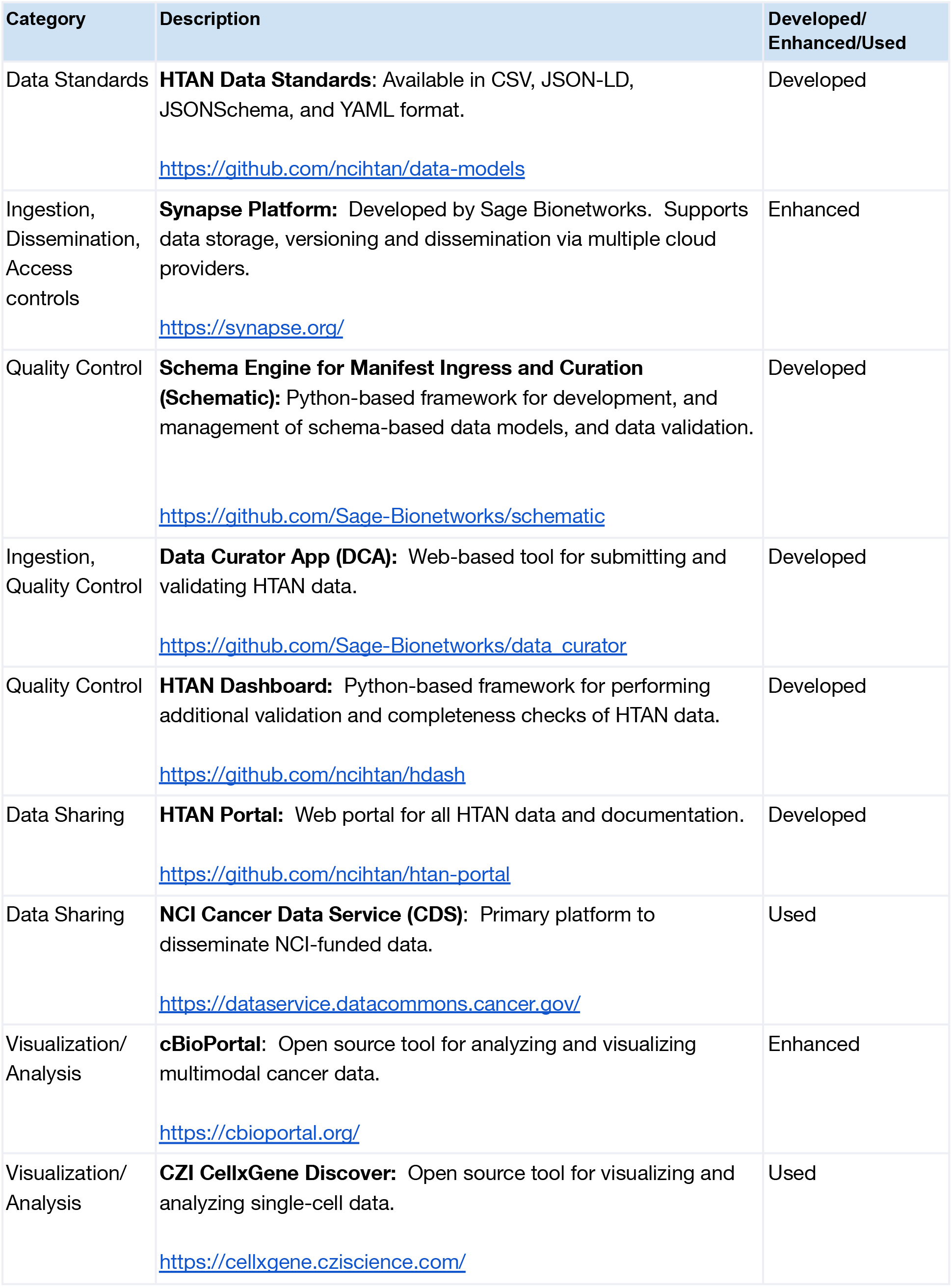
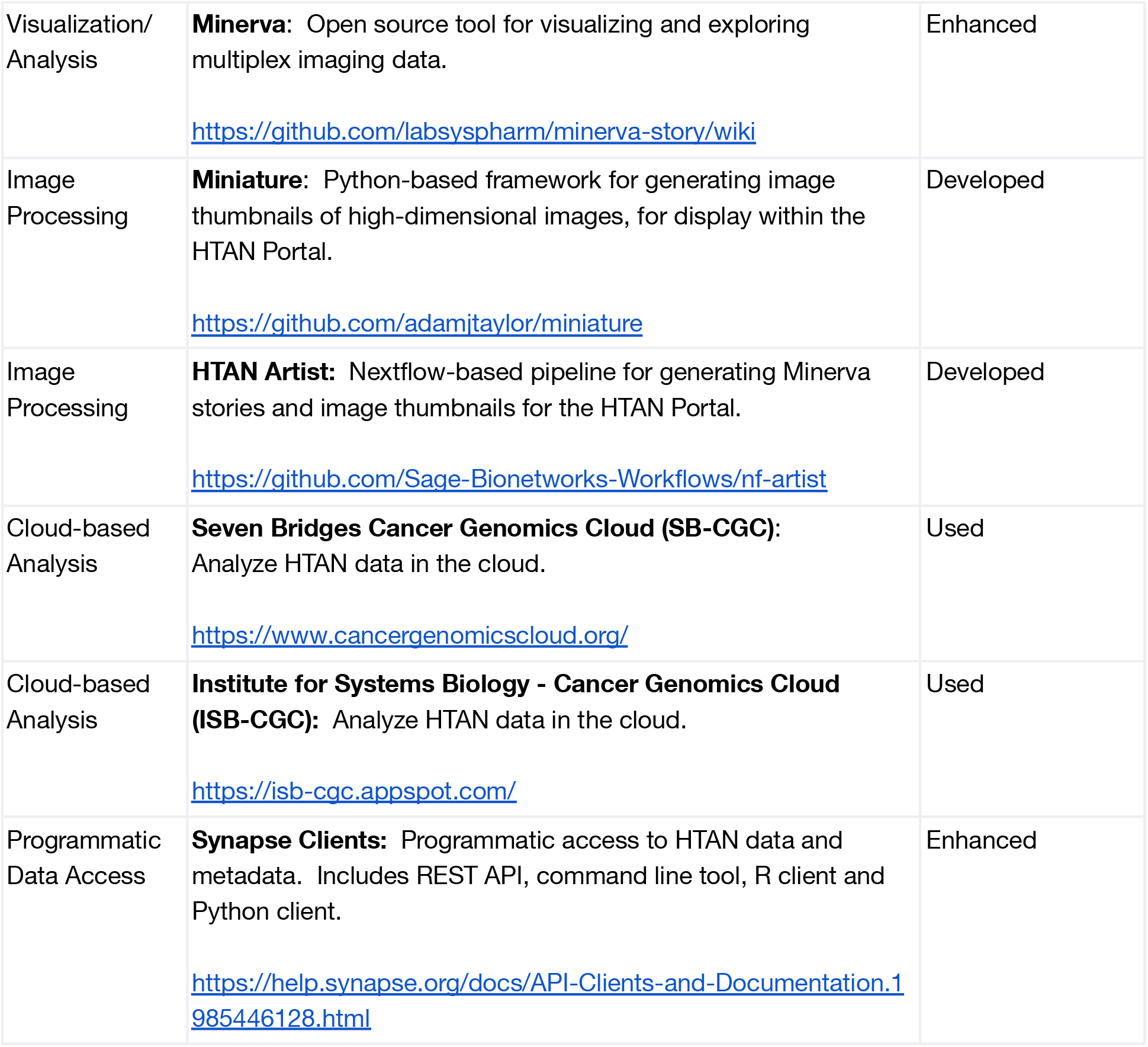
Major data standards, tools and platforms developed, enhanced or leveraged to support HTAN data infrastructure.

| Category | Description | Developed/<br>Enhanced/Used |
| --- | --- | --- |
| Data Standards | **HTAN Data Standards:** Available in CSV, JSON-LD, JSONSchema, and YAML format.<br>https://github.com/ncihtan/data-models | Developed |
| Ingestion, Dissemination, Access controls | **Synapse Platform:** Developed by Sage Bionetworks. Supports data storage, versioning and dissemination via multiple cloud providers.<br>https://synapse.org/ | Enhanced |
| Quality Control | **Schema Engine for Manifest Ingress and Curation (Schematic):** Python-based framework for development, and management of schema-based data models, and data validation.<br>https://github.com/Sage-Bionetworks/schematic | Developed |
| Ingestion, Quality Control | **Data Curator App (DCA):** Web-based tool for submitting and validating HTAN data.<br>https://github.com/Sage-Bionetworks/data_curator | Developed |
| Quality Control | **HTAN Dashboard:** Python-based framework for performing additional validation and completeness checks of HTAN data.<br>https://github.com/ncihtan/hdash | Developed |
| Data Sharing | **HTAN Portal:** Web portal for all HTAN data and documentation.<br>https://github.com/ncihtan/htan-portal | Developed |
| Data Sharing | **NCI Cancer Data Service (CDS):** Primary platform to disseminate NCI-funded data.<br>https://dataservice.datacommons.cancer.gov/ | Used |
| Visualization/ Analysis | **cBioPortal:** Open source tool for analyzing and visualizing multimodal cancer data.<br>https://cbioportal.org/ | Enhanced |
| Visualization/ Analysis | **CZI CellxGene Discover:** Open source tool for visualizing and analyzing single-cell data.<br>https://cellxgene.cziscience.com/ | Used |
| Visualization/<br>Analysis | **Minerva:** Open source tool for visualizing and exploring multiplex imaging data.<br>https://github.com/labsyspharm/minerva-story/wiki | Enhanced |
| Image<br>Processing | **Miniature:** Python-based framework for generating image thumbnails of high-dimensional images, for display within the HTAN Portal.<br>https://github.com/adamjtaylor/miniature | Developed |
| Image<br>Processing | **HTAN Artist:** Nextflow-based pipeline for generating Minerva stories and image thumbnails for the HTAN Portal.<br>https://github.com/Sage-Bionetworks-Workflows/nf-artist | Developed |
| Cloud-based<br>Analysis | **Seven Bridges Cancer Genomics Cloud (SB-CGC):** Analyze HTAN data in the cloud.<br>https://www.cancergenomicscloud.org/ | Used |
| Cloud-based<br>Analysis | **Institute for Systems Biology - Cancer Genomics Cloud (ISB-CGC):** Analyze HTAN data in the cloud.<br>https://isb-cgc.appspot.com/ | Used |
| Programmatic<br>Data Access | **Synapse Clients:** Programmatic access to HTAN data and metadata. Includes REST API, command line tool, R client and Python client.<br>https://help.synapse.org/docs/API-Clients-and-Documentation.1985446128.html | Enhanced |

### Governance and Policy

Responsible data sharing requires transparent governance approaches to ensure data contributors, data curators, and data consumers are empowered to share and use data effectively. To this end, members of the DCC work closely with members of the HTAN consortium to develop and implement data sharing agreements and operational policies following the principles of the NCI Cancer Moonshot Public Access and Data Sharing Policy^19^. A Data and Materials Sharing Agreement (DMSA) establishes the responsibilities and boundaries associated with data sharing with the DCC and within the HTAN consortium, while an External Data Sharing Policy ensures a commitment to share all generated data publicly. An Associate Membership Policy provides a mechanism for experts from outside of HTAN to contribute their expertise and knowledge. In addition, specific policies related to publications and the sharing of research protocols and computational tools were established and implemented. Policy documents can be found on the HTAN Portal. A key infrastructure component for the implementation of these data-sharing policies is the Synapse platform, which provides fine-grained access controls at individual and team levels to ensure that data contributors and DCC staff have appropriate access to their data. Synapse teams were used to enable project-level access and tracked against a table of HTAN membership.

Ensuring the privacy of HTAN research participants is critical and a joint responsibility of the HTAN Centers generating data and the DCC. HTAN Centers are required to fully de-identify data before submission to the DCC via the Synapse platform, and must describe their data and metadata de-identification process in a de-identification plan. Following data submission, the DCC is responsible for implementing additional checks, to ensure patient privacy. Verifying that HTAN imaging data is de-identified is a particular focus of the DCC. For example, the extensive metadata collected alongside HTAN data was noted to theoretically enable the reconstruction of HIPAA-protected dates (such as participant date of birth) from the date of imaging data acquisition through longitudinal metadata attributes. The HTAN DCC therefore developed policies and procedures to confirm that all date attributes, including those in TIFF tags, OME-XML, and other locations, were detected and reported back to data contributors for removal before data release. Throughout, the experience of dedicated governance experts within the DCC and contributions of HTAN members through a policy working group were essential for ensuring alignment and consent across the community.

### Community Engagement

As with any large-scale scientific consortium, it is critical to ensure transparent communication and coordination among principal investigators, data contributors, method and tool developers, as well as other key stakeholders, and to ensure broader engagement with the wider scientific community. Within the consortium, the DCC works to engage all HTAN members at multiple levels of involvement. This includes biannual face-to-face meetings, junior investigator workshops, data workshops, and working groups devoted to policy implementation and scientific collaboration. As noted previously, working groups also drive the RFC process for developing and evolving the HTAN data model.

DCC staff are assigned to both support specific HTAN Centers (i.e., as liaisons) and cover technical areas such as imaging data or clinical metadata. These data liaisons act as named points of contact and facilitate communication between the contributing HTAN Centers and the DCC. Private Slack channels and a help desk ensure data contributors can engage the DCC both for responsive questions and to track bugs or submission issues.

In engaging the wider scientific community, the DCC focuses on timely data releases, outreach to other scientific consortia, and public workshops, e.g., through Data Jamborees and at scientific conferences. We also actively maintain an HTAN manual (https://docs.humantumoratlas.org), our primary external-facing documentation, designed to explain the consortium to new users. The manual describes available HTAN data, HTAN data standards, and all modes of data access. The HTAN help desk is open to external researchers to ask data-specific questions. Finally, we ensure that HTAN data is available via multiple modes of data access across the NCI cancer data ecosystem via the CRDC^5,6^.

## Methods

### HTAN Data Submission Process

The DCC has developed a standardized data submission process (**Fig. 3A**). The process begins with a data curator or scientist from an HTAN Center uploading their data to cloud buckets connected to Synapse. Once the data are uploaded, the submitter needs to provide metadata about each file, including information about its processing and the research participant and biospecimen that it applies to. These metadata are critical for data access and reuse. The metadata is submitted via the Data Curator App (DCA) (**Fig. 3B**), which creates a metadata template based on the data model, validates the provided metadata against the data model, and uploads it to Synapse. Centers also have the option of submitting a filled metadata template describing individual publications and all data associated with a publication.

**Figure 3.**
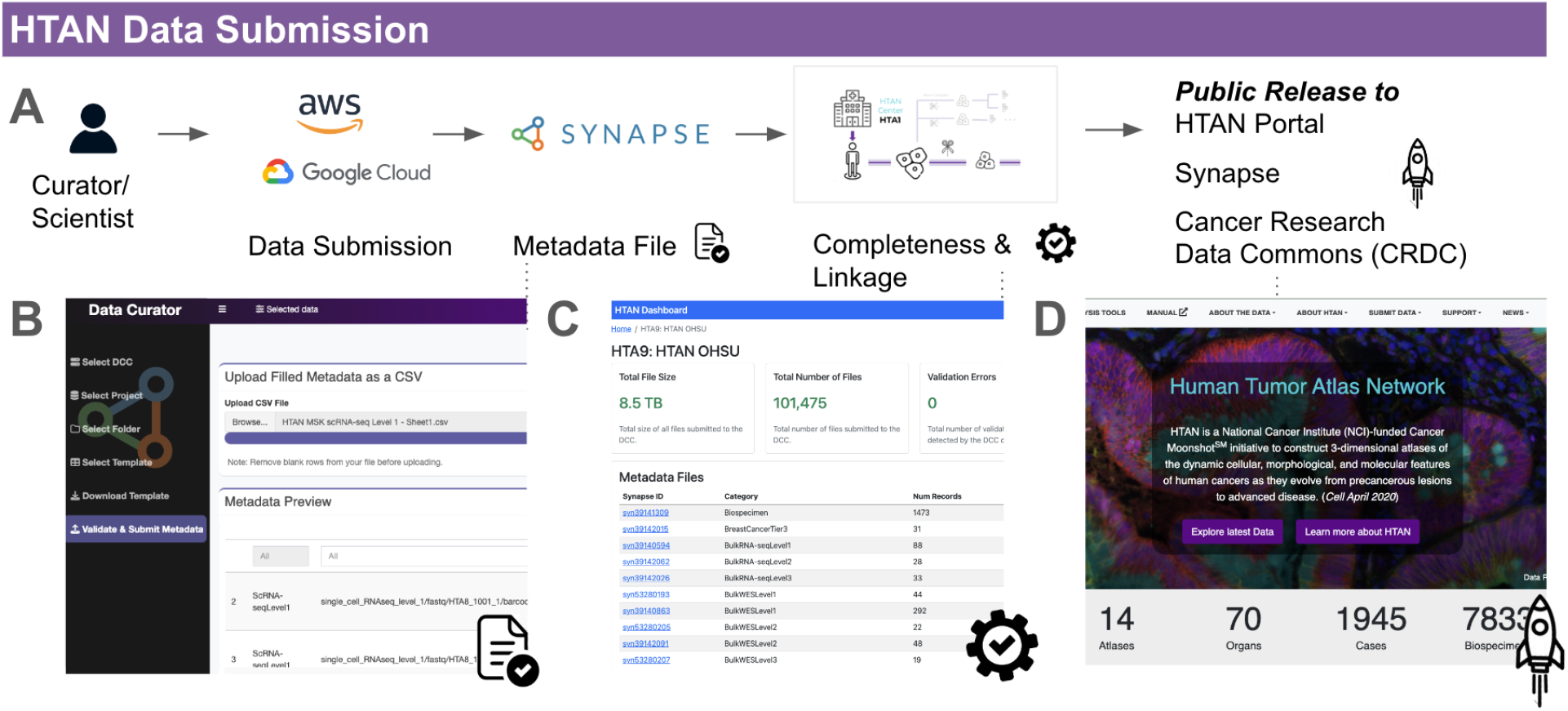
HTAN Data submission and release process. A) An HTAN data curator or scientist uploads data to AWS, Google Cloud, or Synapse, provides metadata about each file, and confirms metadata validation. The DCC performs additional QC checks and releases data to the public. B) the Data Curator App (DCA) performs metadata validation., C) the HTAN Dashboard performs additional QC data checks and checks for overall data completeness. D) the DCC releases the data to the public.

After metadata submission, a second set of validation checks is automatically performed. These checks examine the HTAN Center’s dataset as a whole, verify that all assay data can be linked to parent biospecimens and research participants, and assess data for overall completeness. The results of these checks are made available via the HTAN Dashboard, which is automatically updated every four hours (**Fig. 3C**).

Upon completion of a new data submission, HTAN DCC members review the HTAN Dashboard and relay validation issues to data submitters at the respective HTAN Center. This feedback cycle continues until all validation errors are resolved. Once signed off by the DCC and the Center, all files intended for release are queued. An HTAN Portal preview instance is generated with all data for the next release. After a final manual check is performed, all release data is deployed to the public HTAN Portal. Higher-level processed data is made available publicly on Synapse. Lower-level access-controlled data is submitted to the CRDC^5,6^, where it is made available in subsequent CRDC releases. The data is also submitted in a parallel process to other platforms, including CellxGene^12^, cBioPortal^9–11^, and ISB-CGC^7^, each with its own release cycles. A future goal is to automate the steps of this broader dissemination.

Setting deadlines for major data releases helps to incentivize Centers to submit data in a timely manner. Major releases are completed twice per year, with minor releases in between on an as needed basis. A complete log of data releases is maintained at: https://data.humantumoratlas.org/data-updates. Although HTAN aims to release data upon generation, in practice, we have found that most Centers submit data closer to manuscript submission as incentivized by publishers’ data access requirements and the desire to ensure high quality of data before release.

### Synapse

Sage Bionetworks employs its data management platform, Synapse (RRID:SCR_006307), as the central repository for the HTAN DCC. Each HTAN Center has a dedicated Synapse project, providing a secure environment for uploading, organizing, and annotating data and metadata before public release. Synapse enhances this process through multiple features, including wikis, entity annotations, tabular annotation views for file exploration, and finely tuned access control settings, creating a user- and machine-friendly data management ecosystem.

Project access on Synapse is regulated through team membership, with adjustable permission levels to ensure appropriate access for both data contributors and DCC staff. Moreover, HTAN’s Synapse projects integrate with external storage solutions, such as AWS S3 and Google Cloud Storage, allowing Centers to choose their preferred storage provider, which can minimize egress costs. This is particularly advantageous for contributors who already have data stored with these providers. The platform supports the synchronization of directly added storage objects into Synapse using serverless architectures, e.g., AWS Lambda and Google Cloud Functions. This integration facilitates efficient data uploads via cloud provider clients while maintaining the ease of use associated with Synapse’s web UI, CLI, and language-specific clients in Python and R. For HTAN, the only requirement around folder structure for each Center is that all submissions are grouped into top-level folders categorized by data type, such as scRNA-seq FASTQ files, imaging OME-TIFFs, or demographic information. The exact naming of files is minimally restrictive, as information about the files is captured in the metadata rather than their naming.

### Data Curator App

The Data Curator App (DCA) (**Fig. 3B**), hosted on AWS Fargate, enables data submitters to associate metadata with the submitted assay data files via a wizard-style interface in the browser. The application backend leverages a Python tool, Schematic (https://github.com/Sage-Bionetworks/schematic), to validate the metadata files against the HTAN data standards and submit data to Synapse. Both DCA and Schematic were developed to support multiple data coordination projects at Sage Bionetworks. The separation of UI (DCA) and programmatic schema validation logic (Schematic) simplifies the reuse of these tools across different projects.

In the metadata submission wizard, data contributors select a template (e.g., metadata for clinical demographics or level 1 single-cell RNA sequencing). A Google Sheets link is generated, allowing users to fill out the metadata template directly online using Google Sheets’ functionalities. The Google Sheets template includes checks for the correctness of particular columns. If preferred, the sheet can also be exported as a delimited text file or Excel spreadsheet. Should a specific template be unavailable, a minimal metadata template is used, with the provision to contact a DCC liaison for further guidance. After completing the template, users submit it, and the DCA then leverages Schematic to do an additional check for schema correctness and submits it to Synapse. DCA allows for updating existing metadata as well, accommodating corrections, compliance adjustments, or additions for new files.

### HTAN Dashboard

The HTAN Dashboard (**Fig. 3C**, https://github.com/ncihtan/hdash), is a web application developed to help data submitters across the HTAN Centers and the DCC to track submitted data and associated metadata. For each HTAN Center, the dashboard performs various checks, including tracing and validating all links from files to samples to research participants, ensuring that HTAN IDs follow the specifications and more. It also calculates metadata completeness scores to assess how complete the provided metadata is, in terms of supplied values compared with empty fields. The HTAN Dashboard is written in Python and leverages the Synapse client to programmatically retrieve each Center’s metadata and file counts.

### Image Visualization on the HTAN Portal

HTAN Centers generate imaging data using a broad array of multiplex imaging assays. As of Release 5.1, HTAN has generated imaging data for >3K biospecimens. To enable initial visualization and exploration of these data directly on the HTAN Portal, we deployed narrative guides using Minerva, a lightweight tool suite for interactive viewing and fast sharing of large image data^8^. While extensively curated and interactive guides with manual channel thresholds, waypoints and ROIs can be generated, we implemented an automatic channel thresholding and grouping approach to generate good first defaults, enabling the rapid generation of over 3,700 pre-rendered Minerva stories. Minerva stories are being enhanced with interactive channel selection and embedded metadata. To facilitate recognition and recall of images and tissue features from multiplexed tissue images we developed Miniature, a novel approach for informative and pleasing thumbnail generation from multiplexed tissue images.

### HTAN Data in CZ CellxGene Discover

Single-cell sequencing data is submitted to CZ CellxGene Discover, available at: https://cellxgene.cziscience.com/. The platform enables users to find, explore, visualize, and analyze published datasets. To ensure integration with other single-cell datasets, HTAN data is harmonized to adhere to the CellxGene schema and data format requirements. Critically, this requires that we annotate cell types using terms from the Cell Ontology initiative (CL, https://obofoundry.org/ontology/cl). Annotation is performed by manual mapping of study annotations (cell phenotypes) to the closest CL terms. For example, there is no term for lymphomyeloid primed progenitor-like blasts^20^ and instead selected hematopoietic multipotent progenitor cell (CL_0000837). Precancer and cancer cell mapping posed a challenge, as CL is largely based on normal cells. Cancer cells are annotated with what is hypothesized to be the healthy originating cell type. In cases where no appropriate cell type terms are available, the most relevant parent ontology is used to describe the cell type.

### Integrating with the Cancer Research Data Commons (CRDC)

HTAN data ingress and standardization processes are integrated with the CRDC ecosystem. Multiple CRDC nodes and services support HTAN data download, query, and processing. Specifically, CDS provides access to HTAN controlled-access sequence and imaging files; SB-CGC provides mechanisms to run a variety of processing workflows on HTAN data at CDS; and ISB-CGC contains HTAN tabular metadata and assay data for flexible queries.

HTAN imaging data is available in its original contributed formats, including OME-TIFF and SVS files through the CDS. Preserving a contributor format facilitates both reproducibility of published studies and interoperability with common processing and visualization tools, including processing suites like MCMICRO^21^ and analysis tools such as Napari^22^ and QuPath^23^. A subset of HTAN imaging data has been ingested to the NCI’s Imaging Data Commons^24^ where it has been converted to DICOM^25^ to provide interoperability with other medical imaging datasets and tooling.

The NCI’s cloud resources allow processing of HTAN data on the cloud. For example, SB-CGC^26^ facilitates selection and processing of HTAN single-cell RNA sequence read-level files, image data files, and read-level spatial transcriptomic data. Within ISB-CGC^7^, HTAN data are made available as Google BigQuery tables, allowing flexible SQL query access. More than 850 assay files are queryable through Google BigQuery, encapsulating data from imaging level 4 and single-cell RNA sequencing level 4 assays, collectively spanning more than 200 million cells across spatial and single-cell datasets. Computational notebooks are provided to illustrate cloud-based querying and processing of HTAN data.

## Conclusion

HTAN is planned to continue for at least another 5 years. We have developed a streamlined infrastructure to ingest and disseminate data and can extend the data model with novel assays as needed. We believe that the model employed here will be useful for data coordinating centers of other consortia and have already seen aspects of it reused in other recently formed consortiums, including The Gray Foundation BRCA Pre-Cancer Atlas and the Break Through Cancer (BTC) Foundation, as well as in other data repositories including the CRDC nodes. As there is a wealth of HTAN data available now, we plan to make the broader community aware via tutorials, webinars, and data jamborees, and streamline the reuse of HTAN data based on user feedback. Our infrastructure roadmap includes improvements to the data ingestion process (e.g., additional data integrity checking), improvements to the data release process (increased automation and improved data tracking), improvements to the data portal (enhanced publication pages), and broader data dissemination, including streamlined releases to CRDC, CellxGene, and other repositories.

## Code Availability

All data standards and most tools are available via GitHub (https://github.com/ncihtan). A detailed list of tooling and corresponding repositories is provided in (**Table 3**).

## Acknowledgments

The HTAN Data Coordinating Center (DCC) is supported by NCI Grants: 1U24CA233243-01 and 3U24CA233243-01S2. Support for this work was provided to Memorial Sloan Kettering Cancer Center by a core grant from the National Cancer Institute (P30 CA008748). We thank all research participants for their contributions, and the HTAN Centers for collecting and providing the data. We acknowledge the contributions of DCC staff past and present, including Adam Abeshouse, Elyse Kozlowski, Liz Williams, Jason Hwee, David Gutman, Sheila Reynolds, Justin Guinney, Priti Kumari, Xengie Doan, and Yooree Chae.

## Author Information

### Contributions

I.D.B., M.N are co-first authors; E.C., J.A.E. contributed equally; I.D.B., M.N., C.L., D.L.G, E.M., D.P., N.S., A.J.T., E.C., V. T. developed data standards; E.M., M.D.F., A.J.T, S.O.S., C.L, X.L., A.L., V.T., E.C. developed DCC infrastructure; I.D.B, M.N., C.L., D.L.G., D.P., A.L., J.A., K.A., X.L., S.N., N.S., A.J.T., V.T., E.C. served as data liaisons: J.M. S.S. P.K.S., A.J.T.: developed tooling for imaging data; C.S. developed data sharing and governance policies; I.D.B, M.N., C.L., A.C., A.L., J.A., K.A., X.L., S.N., S.V., N.S., A.J.T., V.T., E.C., J.A.E. wrote the manuscript; A.C., A.L.: coordinated projects, meetings, and communication; N.S., A.J.T., V.T., E.C., J.A.E supervised the HTAN DCC.

## Corresponding authors

Correspondence to Ino de Bruijn or Ethan Cerami.

## Ethics Declaration

### Competing Interests

P.K.S. is a cofounder and member of the BOD of Glencoe Software, member of the BOD for Applied BioMath and a member of the SAB for RareCyte, NanoString, Reverb Therapeutics and Montai Health; he holds equity in Glencoe, Applied BioMath and RareCyte. S.S. is a consultant for RareCyte Inc. Other authors declare no competing interests

## Notes

https://humantumoratlas.org/

## Bibliography

1. Sharpless, N. E. & Singer, D. S. Progress and potential: The Cancer Moonshot. Cancer Cell 39, 889–894 (2021).

2. Hutter, C. & Zenklusen, J. C. The Cancer Genome Atlas: Creating Lasting Value beyond Its Data. Cell 173, 283–285 (2018).

3. Regev, A. et al. The human cell atlas. eLife 6, (2017).

4. Jain, S. et al. Advances and prospects for the Human BioMolecular Atlas Program (HuBMAP). Nat. Cell Biol. 25, 1089–1100 (2023).

5. Hinkson, I. V. et al. A comprehensive infrastructure for big data in cancer research: accelerating cancer research and precision medicine. Front. Cell Dev. Biol. 5, 83 (2017).

6. Wang, Z. et al. NCI cancer research data commons: resources to share key cancer data. Cancer Res. 84, 1388–1395 (2024).

7. Reynolds, S. M. et al. The ISB Cancer Genomics Cloud: A Flexible Cloud-Based Platform for Cancer Genomics Research. Cancer Res. 77, e7–e10 (2017).

8. Hoffer, J. et al. Minerva: a light-weight, narrative image browser for multiplexed tissue images. J. Open Source Softw. 5, (2020).

9. Cerami, E. et al. The cBio cancer genomics portal: an open platform for exploring multidimensional cancer genomics data. Cancer Discov. 2, 401–404 (2012).

10. Gao, J. et al. Integrative analysis of complex cancer genomics and clinical profiles using the cBioPortal. Sci. Signal. 6, pl1 (2013).

11. de Bruijn, I. et al. Analysis and Visualization of Longitudinal Genomic and Clinical Data from the AACR Project GENIE Biopharma Collaborative in cBioPortal. Cancer Res. 83, 3861–3867 (2023).

12. Megill, C. et al. cellxgene: a performant, scalable exploration platform for high dimensional sparse matrices. BioRxiv (2021) doi:10.1101/2021.04.05.438318.

13. CZI Single-Cell Biology et al. CZ CELLxGENE Discover: A single-cell data platform for scalable exploration, analysis and modeling of aggregated data. BioRxiv (2023) doi:10.1101/2023.10.30.563174.

14. Thorogood, A. et al. International federation of genomic medicine databases using GA4GH standards. Cell Genomics 1, (2021).

15. Heath, A. P. et al. The NCI genomic data commons. Nat. Genet. 53, 257–262 (2021).

16. Schapiro, D. et al. MITI minimum information guidelines for highly multiplexed tissue images. Nat. Methods 19, 262–267 (2022).

17. Benjelloun, O., Chen, S. & Noy, N. Google Dataset Search by the Numbers. (2020).

18. Warzel, D. B. et al. Common data element (CDE) management and deployment in clinical trials. AMIA Annu. Symp. Proc. 1048 (2003).

19. Cancer MoonshotSM Public Access and Data Sharing Policy - NCI. https://www.cancer.gov/research/key-initiatives/moonshot-cancer-initiative/funding/public-access-policy.

20. Chen, C. et al. Single-cell multiomics reveals increased plasticity, resistant populations, and stem-cell-like blasts in KMT2A-rearranged leukemia. Blood 139, 2198–2211 (2022).

21. Schapiro, D. et al. MCMICRO: a scalable, modular image-processing pipeline for multiplexed tissue imaging. Nat. Methods 19, 311–315 (2022).

22. Napari Contributors. napari: a multi-dimensional image viewer for python. Zenodo (2019) doi:10.5281/zenodo.3555620.

23. Bankhead, P. et al. QuPath: Open source software for digital pathology image analysis. Sci. Rep. 7, 16878 (2017).

24. Fedorov, A. et al. NCI imaging data commons. Cancer Res. 81, 4188–4193 (2021).

25. National Electrical Manufacturers Association. NEMA PS3 / ISO 12052 Digital Imaging and Communications in Medicine (DICOM) Standard. https://www.dicomstandard.org/.

26. Lau, J. W. et al. The Cancer Genomics Cloud: Collaborative, Reproducible, and Democratized-A New Paradigm in Large-Scale Computational Research. Cancer Res. 77, e3–e6 (2017).

